# AI-Designed, Mutation-Resistant Broad Neutralizing Antibodies Against Multiple SARS-CoV-2 Strains

**DOI:** 10.1101/2023.03.25.534209

**Authors:** Yue Kang, Yang Jiao, Kevin Jin, Lurong Pan

## Abstract

In this study, we generated a Digital Twin for SARS-CoV-2 by integrating data and meta-data with multiple data types and processing strategies, including machine learning, natural language processing, protein structural modeling, and protein sequence language modeling. This approach enabled the computational design of broadly neutralizing antibodies against over 1300 different historical strains of SARS-COV-2 containing 64 mutations in the receptor binding domain (RBD) region. The AI-designed antibodies were experimentally validated in real-virus neutralization assays against multiple strains including the newer Omicron strains that were not included in the initial design base. Many of these antibodies demonstrate strong binding capability in ELISA assays against the RBD of multiple strains. These results could help shape future therapeutic design for existing strains, as well as predicting hidden patterns in viral evolution that can be learned by AI for developing future antiviral treatments.

## Introduction

The evolving strains of the SARS-CoV-29 virus have rendered many previously approved antibody therapeutics ineffective, especially if they target the Spike protein of the virus [REF]. Among the hundreds of mutation variations observed along its phylogenetic tree, mutations within the ACE2 Receptor binding domain (RBD) of the Spike protein have been the major focus for researchers, as these mutations drastically affect the binding strength of Spike with the ACE2 receptor. For instance, the L452R substitution found in the B.1.427 and B.1.429 lineages leads to a significant reduction in susceptibility to bamlanivimab [1], as well as a modest decrease in susceptibility to the combination of bamlanivimab and etesevimab [1,2,3].

In the present study, we leveraged artificial intelligence (AI) to perform *in silico* mutagenesis and screening on more than 10^9^ antibody sequences in order to identify candidates that bind broadly with known previously observed Spike protein RBD variants with high affinities. Graph neural networks (GNNs) [8] are neural network architectures designed specifically to cope with graph data. Nodes in the graph are designed to learn an embedding which contains information about their associated neighbors. The embeddings can function as characteristic features for node labeling, edge prediction, and graph representation with proper readout and pooling methods [9, 10, 11, 12]. The intrinsic design of GNN makes it well-suited to study molecular and biological interactions and other chemical and physical properties. We therefore seek to describe antibody-antigen interactions in a graph-based manner.

Language-based networks can also model proteins given the assumption that the primary protein structure is analogous to natural language sequences [13, 14, 15, 16, 17, 18]. Hidden dependencies and interactions between amino acids may be trainable by the temporal dynamic inherently designed in the basic recurrent neural networks, such as long short-term memory network (LSTM) and Transformer neural networks.

We explored various modeling strategies from these domains, including graph neural networks (GNN) and natural language processing (NLP) architectures, in which protein sequences are described with a graph-based representation and a language-based representation, respectively.

We also hypothesized that the cross-binding antibodies may possess mutation-resistant binding capability towards future evolving RBD region variations, and therefore may demonstrate broad neutralization potency, a key characteristic of therapeutic antibodies targeting rapidly mutating viruses such as SARS-CoV-2. In addition, the efficient nature of this AI-driven antibody discovery approach lays the groundwork for future fast response therapeutic discovery in pandemic preparedness. This approach also opens new doors for conventional protein drug discovery in targeting multiple antigens for selectivity optimization or broad binding.

## Methods

### 1. In-silico antibody affinity maturation modeling via AI

#### 1.1 Training and testing datasets

Ainnocence has developed antibody affinity maturation AI models based on deep neural networks. Publicly available curated datasets, including the SKEMPI database[4] Antibody-Bind (AB-Bind) database [5], Observed Antibody Space[6], and UniProt[7] were examined during model development. The resulting model’s performance was validated on SKEMPI and AB-Bind, and both datasets curated a protein-protein complex with relevant PDB structures, along with single-site and multi-site mutations. In AB-Bind, the binding affinities for mutated variants are represented by the change in free energy (∆∆G) of binding in kcal/mol, whereas in SKEMPI, both the affinities of wild-type complexes and the affinities of the mutants are provided with experimental dissociation constant (Kd) values.

#### 1.2 Deep Learning models

To reduce computing cost and extend the model application to proteins, the structures of which have not been solved, we explored a model that required only a protein’s amino acid sequence as input to predict binding affinity. The proposed study aims to examine whether antibody-antigen complex affinity prediction can be accomplished using primary sequence inputs only. Figure 1 illustrates several examples of modeling strategies.

**Figure 1:**
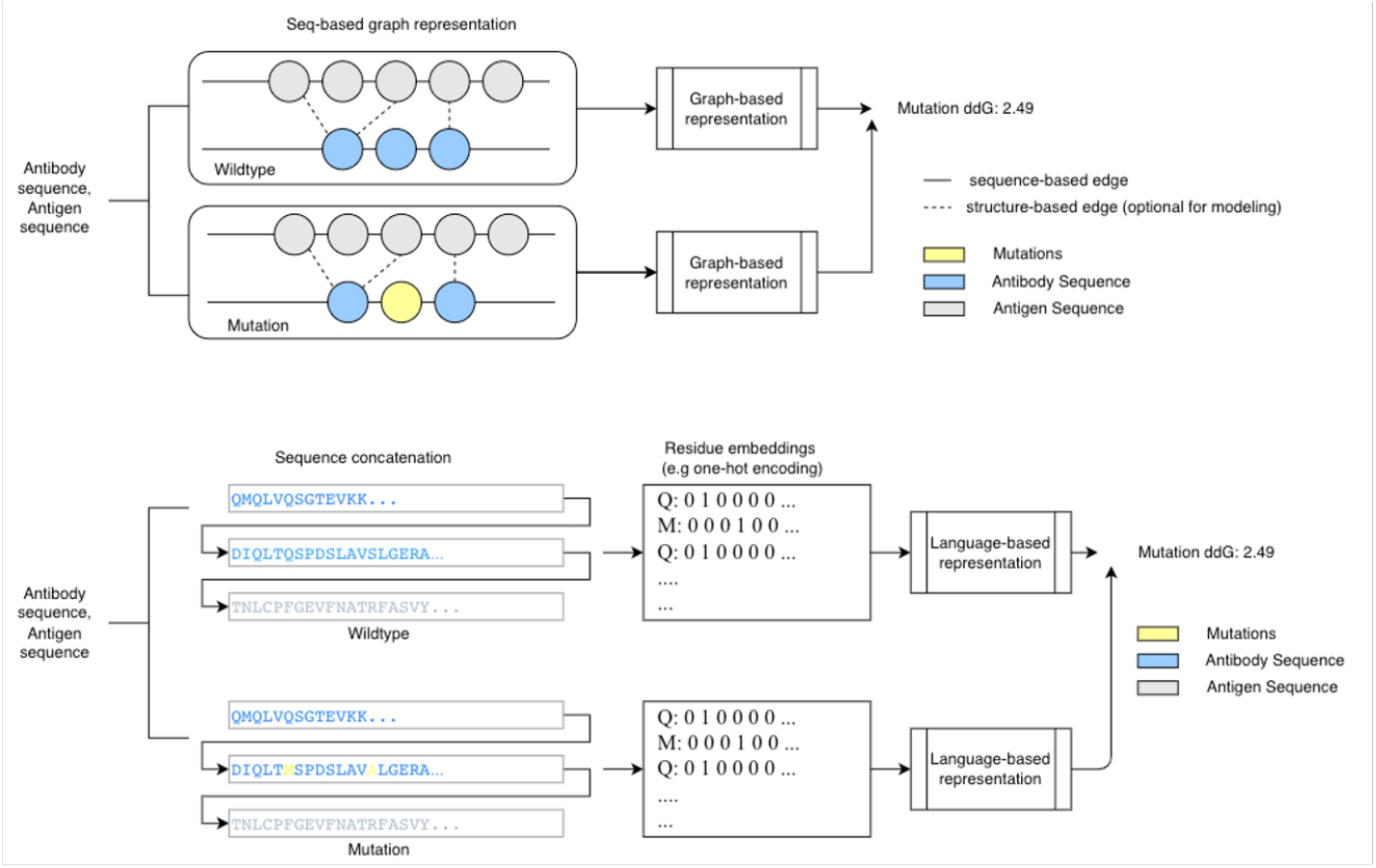
Design of the two modeling approaches developed to support sequence-based antibody affinity design. (A) Graph neural network representation for affinity training (B) Language-based (BiLSTM) neural network representation for affinity training

The prediction of binding affinity variations between mutated antibodies and a given antigens is accomplished by classification tasks. We consider mutations that result in increased binding affinity (i.e. ∆∆G <0) as positive samples, and mutations that result in decreased binding affinity (i.e. ∆∆G >0) as negative samples. Our model seeks to extract latent properties of amino acids that can be best used for optimal delineation between ‘strengthened’ and ‘weakened’ binders.

#### 1.3 Model Evaluation

We performed out-of-distribution cross-validation to evaluate the model’s accuracy as well as scalability. The resulting trained models were later summarized and engineered and function as the antibody maturation engine for SentinusAI™, our purpose-built structure-free large molecule platform.

In practice, the *in silico* affinity maturation of SentinusAI™ mimics the natural immune system to search for high affinity binders towards the given target/antigen within a virtual library of size up to 10^15^, allowing it to cover a somatic mutation space that is not feasible in a wet-lab environment.

Later, a 10,000-times faster efficiency compared to traditional methods was also achieved with optimized engineering and distributed computation strategy.

### 2. Computational workflow for identifying COVID-19 neutralizing antibodies

#### 2.1 Data Collection

Information of over 1300 different historical SARS-CoV-2 strains (including wildtype (B.1) and Delta) were retrieved from the GISAID [19, 20, 21] database as of August 26, 2021. Three wildtype antibodies, CR3022, Casirivimab (Regen 10933), and Imdevimab (Regen 10987) were chosen as templates for in -silico cross-binding antibody design. The CR3022 antibody [22] was a SARS-CoV-1-specific monoclonal antibody obtained from human convalescent plasma in a patient that had recovered from severe acute respiratory syndrome (SARS-CoV-1), a virus closely related to the novel coronavirus that causes COVID-19. CR3022 cross-reacts with the novel coronavirus, although the antibody does not bind tightly enough to neutralize and stop it from infecting cells [22, 23, 24, 25]. Both Casirivimab and Imdevimab [26], are monoclonal antibody cocktails developed by Regeneron with validated potency on wildtype as well as Delta strain.

#### 2.2 *In silico* mutant library generation

To generate an in silico mutant library, we only consider mutations on the template antibody paratope. In order to map the antibody paratope, we obtained the crystal structure of the SARS-CoV-2 RBD in complex with template antibody CR3022 from the PDB bank [PDB ID: 6W41]. Interface contacts were identified within the 5.5 Å distance cutoff. Single-site and double-site mutations were then exhaustively generated within the paratope on both heavy and light chains to form an antibody somatic mutation library (Figure 2: Step 1). Using this strategy, more than 109 mutants were derived from each antibody template (CR3022, Regen 10933, and Regen 10987).

**Figure 2:**
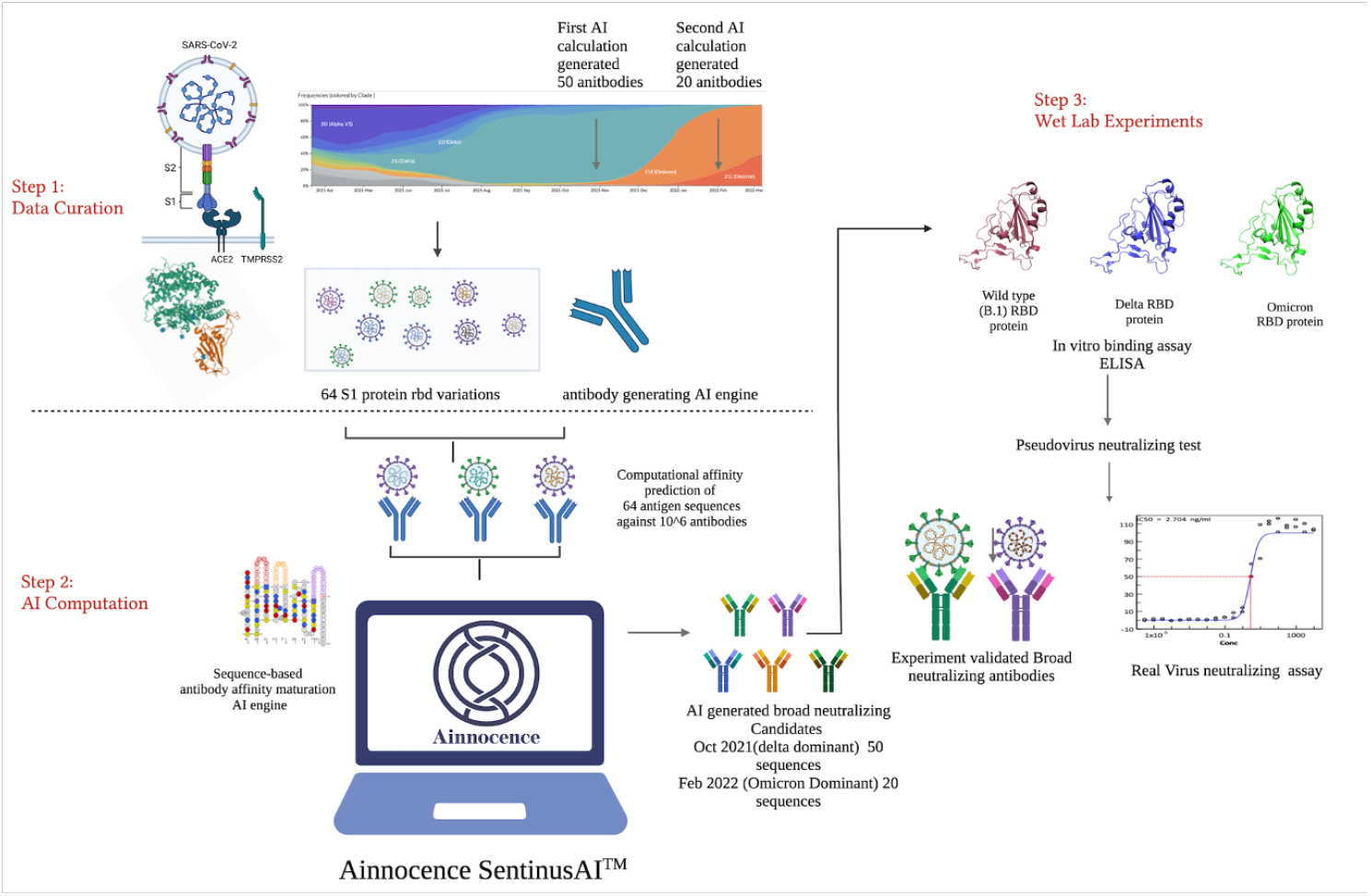
AI-based broad-neutralizing antibody design of SARS-COV-2. **Step1:** SARS-CoV-2 Cross-binding sequence selection and virus mutation data curation. **Step2:** AI-based antibody binding prediction and cross-variants binding selection for potential candidate sequences for future variants **Step3:** ELISA-based assay for antibody’s binding ability measurement; Neutralization assay and cytopathic effect (CPE) reduction assay for antibody’s neutralizing capacity measurement.

#### 2.3 *In silico* library generation

During our first round of in silico antibody design, the goal was to discover antibodies with improved binding affinities with over 1300 different historical SARS-CoV-2 strains, including wildtype (B.1) strain and Delta strain. Therefore, antibody sequences with high cross-binding scores were selected. For each mutant antibody in the mutagenesis library and each unique mutant S1 protein of the historical SARS-CoV-2 strains, their affinity scores were computed based on their VH and VL chain and viral S1 protein sequences as the antigen. This score represents the likelihood of affinity improvement of a given antibody towards the targeting antigen. For each SARS-CoV-2 spike protein mutant, we selected the top 200 highest-scored antibodies from the 109 somatic mutation space as ‘strong binders’ for that specific strain. All ‘strong binders’ were then computed and selected following the above protocol for all 1300 SARS-CoV-2 strains (including B.1 and Delta) as described in Step 2 of Figure 2.

SARS-CoV-2 variants contain mutations in different regions of the S1 protein along the evolutionary tree, so we hypothesized that antibodies that could bind to all observed S1 protein may be able to bind future variants with S1 muta During our first round of in silico antibody design, the goal was to discover antibodies with improved binding affinities with over 1300 different historical SARS-CoV-2 strains, including wildtype (B.1) strain and Delta strain. Therefore, antibody sequences with high cross-binding scores were selected. For each mutant antibody in the mutagenesis library and each unique mutant S1 protein of the historical SARS-CoV-2 strains, their affinity scores were computed based on their VH and VL chain and viral S1 protein sequences as the antigen. This score represents the likelihood of affinity improvement of a given antibody towards the targeting antigen. For each SARS-CoV-2 spike protein mutant, we selected the top 200 highest-scored antibodies from the 109 somatic mutation space as ‘strong binders’ for that specific strain. All ‘strong binders’ were then computed and selected following the above protocol for all 1300 SARS-CoV-2 strains (including B.1 and Delta) as described in Step 2 of Figure 2.

SARS-CoV-2 variants contain mutations in different regions of the S1 protein along the evolutionary tree, so we hypothesized that antibodies that could bind to all observed S1 protein may be able to bind future variants with S1 mutations. From the common strong-binding antibody candidates for all 1300 variants, we selected the top 50 cross-binding candidates which have the highest average predicted score among all variants. This concludes our first-round computations for cross-binding antibodies prior to Omicron.

A second round of computation was performed in February 2022 to further improve the Ab affinity towards Omicron. The same procedures detailed above were followed and the top 20 cross-binding antibody sequences with the highest average prediction scores were selected. From the common strong-binding antibody candidates for all 1300 variants, we selected the top 50 cross-binding candidates which have the highest average predicted score among all variants. This concludes our first-round computations for cross-binding antibodies prior to Omicron. Overall, this round further improve the Ab affinity towards Omicron. The same procedures detailed above were followed and the top 20 cross-binding antibody sequences with the highest average prediction scores were selected.

### 3. Wet lab experimentation

#### 3.1 ELISA assays

An ELISA-based assay was used to measure the antibody’s ability to bind to the RBD of the different SARS-CoV-2 strains (B.1, Delta, Omicron).

To prepare the ELISA plate, coating buffer (500 mL) containing CBS (0.75 g) and NaCO_3_/NaHCO_3_ (1.46 g) was adjusted to pH = 9.6. Each of the RBD proteins of three virus strains (B.1, Delta, Omicron) were prepared at concentrations of 0.03 μg/ml and 1 μg/ml., 100 μl was added to each well, and coated at 4 °C overnight. The solution in the plate was removed by shaking and panting, and 2% of BSA was added to each well for sealing. The proteins were incubated for 1 hour at room temperature. The solution was then discarded, and the proteins were washed twice with 300 μl of elution buffer (PBS buffer containing Tween-20 0.2%, pH = 7.2 -7.4) and patted dry.

Each antibody was diluted to 1 μg/ml with the elution buffer containing 0.1% BSA., 100 μl of diluted antibody was added to each well previously coated with RBD proteins, mixed evenly and incubated at room temperature for 2 h. The solution was discarded, and the antibody was washed three times with 300 μl of elution buffer and patted dry. 100 μl of Jackson: Goat Anti-Human IgG (H+L)/HRP secondary antibody at a working concentration was added to each well, mixed evenly, and incubated at room temperature for 1 h. The mixed solution was discarded, and the antibody was washed with 300 μl of elution buffer and patted dry. The TMB substrate solutions A and B were mixed evenly at a ratio of 1:1. Then, 200 μl of solution mixture was added to each well and incubated at room temperature in a dark room; 50 μl of 2M sulfuric acid stop solution was added to each well, and an OD value was measured immediately at 450 nm. Overall, ELISA data were generated from two different concentrations (0.03 ug/ml, 1 ug/ml) of RBD (WT, Delta, Omicron) proteins and 7 serial dilutions of the generated monoclonal antibodies.

#### 3.2 Pseudo virus-based neutralization assay

The pseudovirus consists of a lipid envelope expressing specific glycoproteins (e.g., from SARS-CoV-2) and a surrogate virus core. Generally, the virus core was genetically modified so that it was unable to express the surface protein on its own. Pseudoviruses are capable of infecting susceptible cells from different species with higher titer and resistance to serum complement, but they only replicate for 1 round in the infected host cells. These well-established pseudovirus assays are therefore safer and easier to perform for virus neutralization assays, particularly for highly infectious and pathogenic viruses such as Covid-19, SARS, Ebola, and H5N1.

First, the SARS-CoV-2 (2019-nCoV) Spike pseudo virus was constructed by Sino Biological, Inc. The pseudo virus consists of a human immunodeficiency virus Type I (HIV-1) backbone, the fragment encoding for the surface SARS-CoV-2 (2019-nCoV) Spike protein, and a luciferase reporter gene for marking infected cells. This pseudo virus could not self-replicate and showed robust infectivity according to the vendor’s published data. The host cell used for transfection is H293T.

Next, for the neutralization assay workflow, we pre-incubated 293T-ACE2 cells with neutralizing antibodies, followed by pseudovirus addition, followed by chemiluminescence detection of luciferase relative light units (RLUs). The antibodies were serially diluted in a 96-well plate (Table 5), followed by pseudovirus addition. The plate was incubated for 1 h at 37 °C so that the pseudovirus bound to the antibodies, followed by addition to pre-plated 293T-ACE2 cells, with duplicate wells for each concentration.

After 48-72 h in culture in a 5% CO2 incubator at 37°C, a luciferase substrate was added and the RLU was assessed. The neutralizing ability of the antibody was calculated by the following equation:

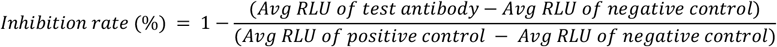

The IC50 values were calculated by the Reed-Muench method.

#### 3.3 Coronavirus cytotoxicity assay

Viruses hijack cellular mechanisms for viral replication, leading to cell death, but antiviral drugs can block this effect. We used a cytopathic effect (CPE) reduction assay to determine whether neutralizing antibodies improved cell viability. This assay was performed with vero E6 cells expressing the ACE2 receptor, which mediates viral infection:

First, cells were cultured in a minimum essential medium (MEM) supplemented with 10% heat-inactivated fetal bovine serum (HI FBS). For the CPE and toxicity assays, cells were collected from MEM supplemented with 1% penicillin streptomycin glutamine (PSG) and 2% HI FBS and resuspended to 200,000 cells per ml. A total of 20 μl cell suspension (about 4000 cells) was added to each well.

Neutralization of the virus was detected by mixing a fixed number of virus particles with serial dilutions of the antibody, followed by the CPE assay in cells. In 384-well plates, we added 5 μl of serum-diluted antibody along with 5 μl of virus containing 1000 TCID, and the plate was incubated for 1 h at 37 °C. The CPE assay was initiated by adding 20 μl of the cell suspension. Blank controls only contained cells, and no antibody was added to the virus controls. After incubating the plate at 37 °C/5% CO2/90% humidity for 72 hours, 30 μl of the Promega Cell Titer-Glo Luminescent Cell Viability Assay Kit was added to each well. Luminescence was read using a Perkin Elmer Envision or BMG CLARIOstar microplate reader following incubation at room temperature for 10 min to measure cell viability. Raw data from each well was normalized to the inhibition rate of 100% without antibody and the inhibition rate of 0 for blank controls to calculate % inhibition of CPE using the following formula:

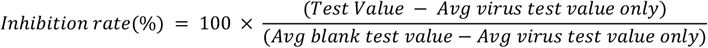

The CPE assay was conducted in a biosafety level-3 laboratory with the plate being sealed with a clear cover prior to reading.

Finally, in the detection of antibody cytotoxicity, antibodies were serially diluted with the same medium as in the CPE assay. A mixture of 20 μl of cells and 10 μl of antibodies was added to each well in a multi-well plate. Cells only served as blank controls, and only cells treated with benzethonium chloride (100 µM final concentration) served as negative controls. Luminescence was generated and read in the same way as in the CPE assay.

## Results

### AI modeling benchmarks

We here investigated different modeling approaches to improve prediction accuracy of antibody maturation based on sequence inputs only. The performance on previously unseen samples has been one of the most common challenges for deep learning neural networks, and our modeling endeavors aim to improve the model’s robustness under the scenarios of antibody affinity predictions. To find an optimized model for antibody screening, several neural network modeling approaches shown in Table 2 (C) were investigated, and their performances were assessed by evaluating the prediction accuracy on previously unseen samples.

The out-of-distribution experiments were designed as follows: five antigens and their associated complexes (wildtype and mutations) were randomly selected as the test set, with the remaining data used as the training set, to make sure that the model never ‘seen’ sequences like the test set during training. We referred to this validation as the ‘leave-5-out’ (L5O) approach.

Performances of classification were assessed by ROC area under curve (AUC). We further evaluate the correlation between experimental affinity changes and predicted values using the Pearson correlation coefficient and Spearman coefficients.

Table 2 (A) shows the averaged AUC of binary classification on improved vs weakened binders using language-based model and graph-based model. Benchmark studies from [5] are listed for comparison.

Both the graph-based (AUC = 0.82) and language-based (AUC = 0.73) modeling approaches were able to distinguish strengthened and weakened binders with better or comparable performance relative to Discovery Studio [5], the best reported structure-based approaches (Table 1). Unlike Discovery studio, which employs a physical model derived from primary, secondary, and tertiary protein structure to compute binding affinity, our model learns the mapping between antibody sequence and binding affinity from a large amount of experimental data.

**Table 1.** Diluted Concentration of Antibody

| S/N | Diluted Concentration<br>(µg/mL) |
| --- | --- |
| 1 | 0.0014 |
| 2 | 0.0041 |
| 3 | 0.012 |
| 4 | 0.037 |
| 5 | 0.11 |
| 6 | 0.33 |
| 7 | 1 |

We then compared the model’s Pearson coefficient with prior works [5] to examine the linear correlation between predicted values and experimental affinity changes upon mutations (Table 2.B, Table 2.C). The graph-based model (Pearson = 0.61) outperformed most conventional (structure-based) *in silico* approaches, whereas language-based prediction yields comparable correlation to Discovery Studio (Pearson = 0.45) with Pearson coefficient of 0.40 (Table 2). These observed correlations between affinity changes and model-based probabilities demonstrates that deep learning-based representations can be utilized in predictions on completely unseen variants during antibody maturation. We hypothesized that the model captures the transferable features that contribute to the binding strength in antibody-antigen interactions. We also compared the two proposed learning-based approaches in terms of their ranking ability, represented by Spearman ranking coefficients (Table 2. C), as well as the model’s prediction performance on several ‘unseen’ complexes (Figure 3).

**Table 2.**
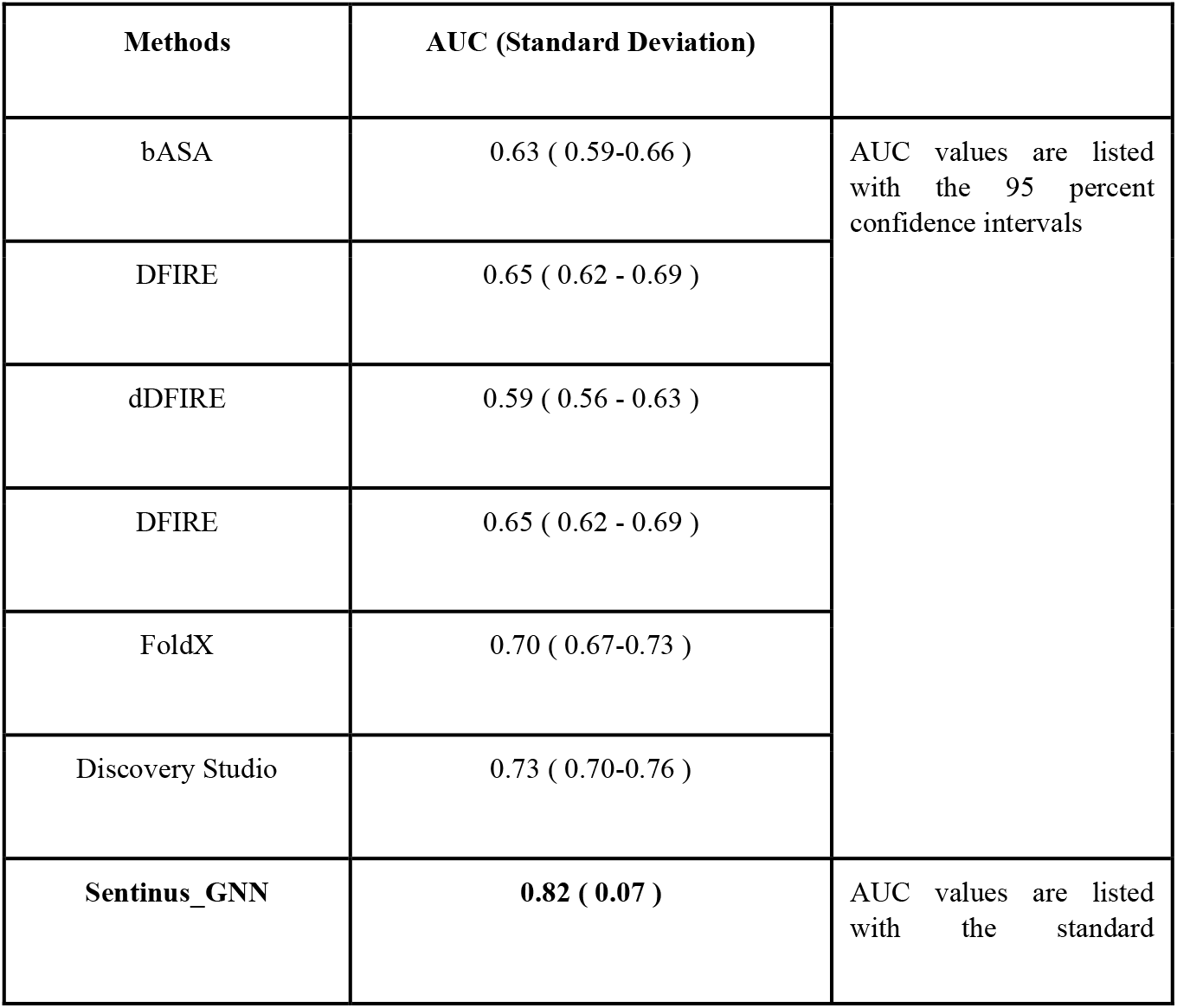
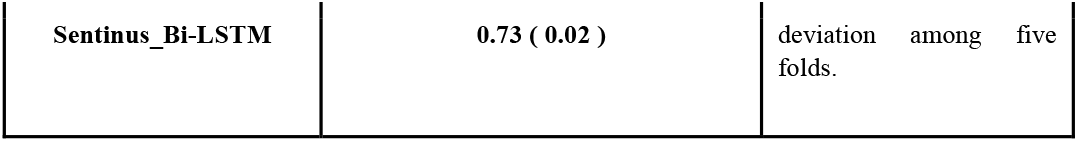
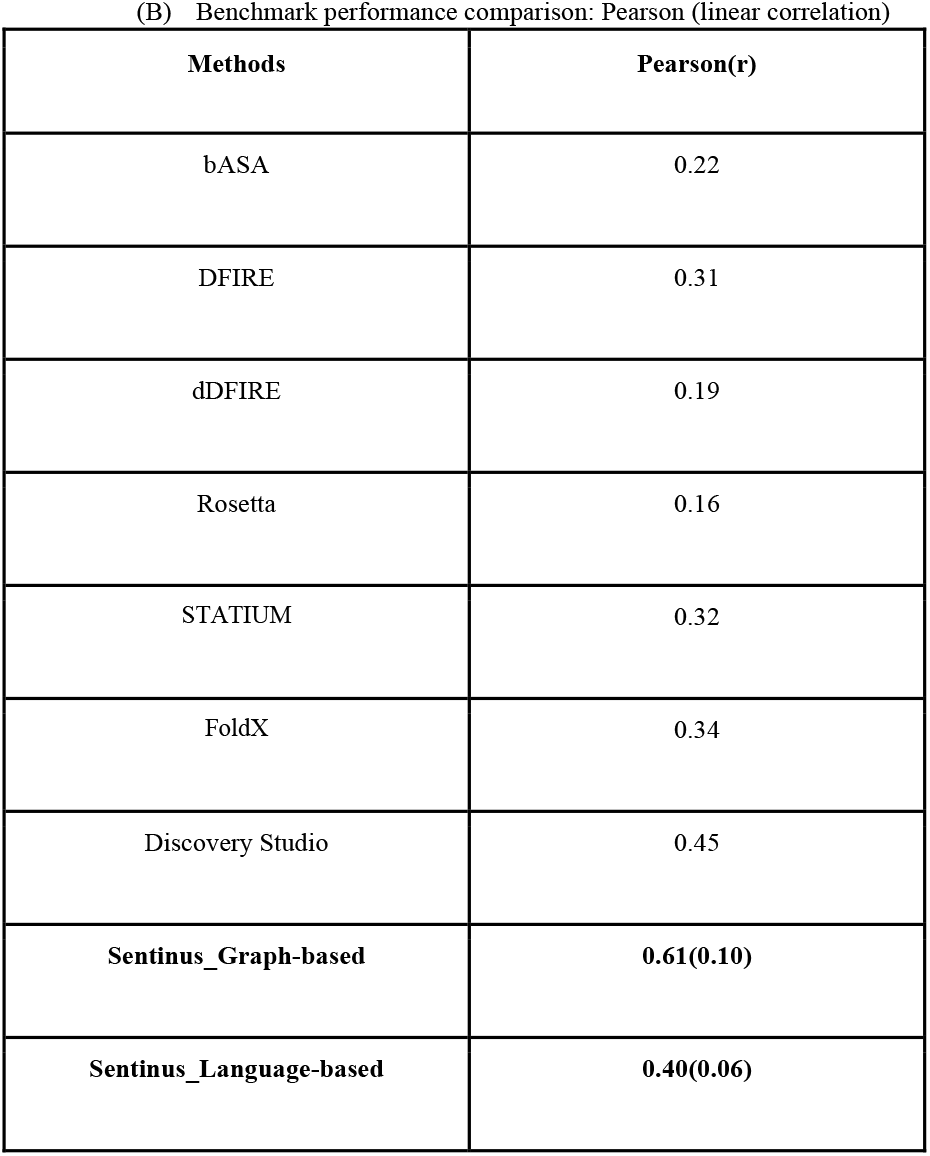
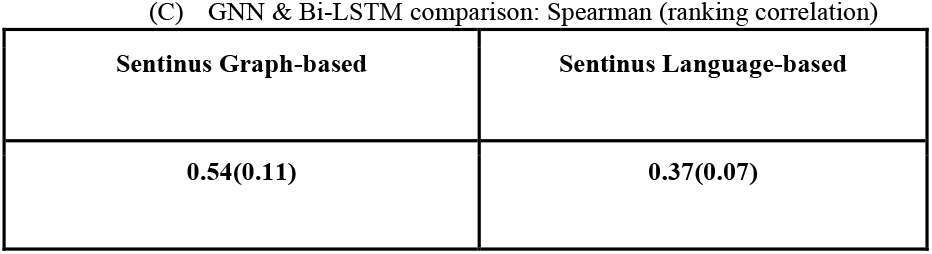
Baseline five-folds out-of-distribution validation

| Methods | AUC (Standard Deviation) |  |
| --- | --- | --- |
| bASA | 0.63 ( 0.59-0.66 ) | AUC values are listed with the 95 percent confidence intervals |
| DFIRE | 0.65 ( 0.62 - 0.69 ) |  |
| dDFIRE | 0.59 ( 0.56 - 0.63 ) |  |
| DFIRE | 0.65 ( 0.62 - 0.69 ) |  |
| FoldX | 0.70 ( 0.67-0.73 ) |  |
| Discovery Studio | 0.73 ( 0.70-0.76 ) |  |
| **Sentinus_GNN** | **0.82 ( 0.07 )** | AUC values are listed with the standard |
| **Sentinus_Bi-LSTM** | **0.73 ( 0.02 )** | deviation among five folds. |

(B) Benchmark performance comparison: Pearson (linear correlation)
| **Methods** | **Pearson(r)** |
| --- | --- |
| bASA | 0.22 |
| DFIRE | 0.31 |
| dDFIRE | 0.19 |
| Rosetta | 0.16 |
| STATIUM | 0.32 |
| FoldX | 0.34 |
| Discovery Studio | 0.45 |
| **Sentinus_Graph-based** | **0.61(0.10)** |
| **Sentinus_Language-based** | **0.40(0.06)** |

Table 2 (A) shows the averaged AUC of binary classification on improved vs weakened binders using language-based model and graph-based model. Benchmark studies from [5] are listed for comparison.
| **Sentinus Graph-based** | **Sentinus Language-based** |
| --- | --- |
| **0.54(0.11)** | **0.37(0.07)** |

**Figure 3:**
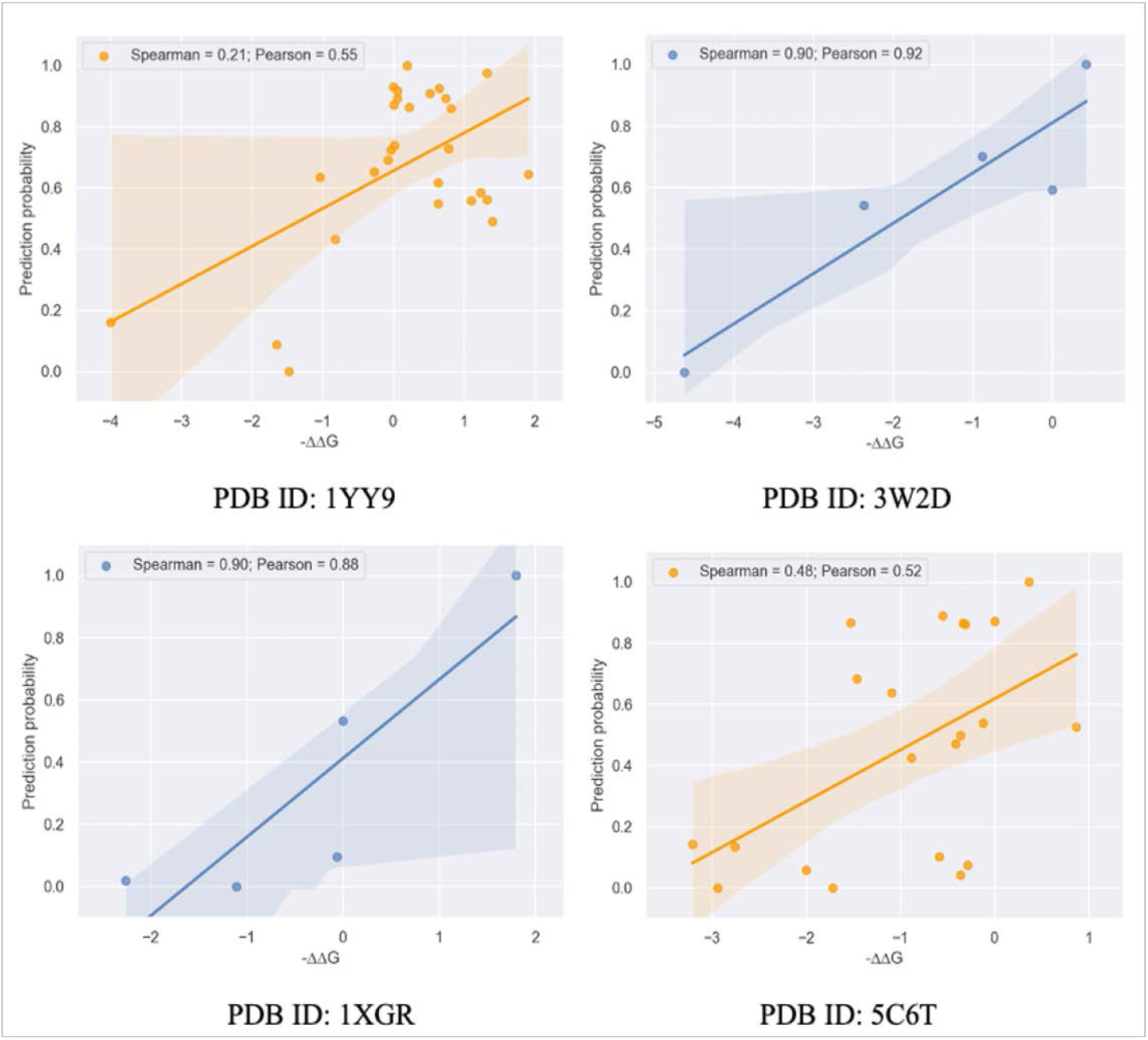
Scatterplot of model-predicted probabilities with experimental ∆∆G values for four representative complexes (wildtypes and mutations) from the out-of-distribution test set. The x-axis represents -∆∆G value, which positively correlates with the affinity strength of each sample, Y-axis represents model predicted probabilities. The correlation can be quantitatively demonstrated by Pearson’s and Spearman’s value respectively.

In summary, both graph-based and natural language-based approaches can be used to predict interactions with novel variants during antibody maturation. We hypothesized that the model captures the key transferable features for predicting binding strength in antibody-antigen interactions.

### Affinity maturation efficiency

The proposed modeling strategy with significantly reduced complexity allows SentinusAI to search large mutation spaces with very low computational cost. Table 3 illustrates the efficiency comparison of in silico affinity maturation with two conventional structure-based approaches. Prodigy (PROtein binDIng enerGY prediction) is a collection of web services focused on the prediction of binding affinity in biological complexes as well as the identification of biological interfaces from crystallography [29]. Schrodinger can also perform molecular dynamic studies to calculate free energy (binding affinity) [28]. Both approaches require complex structural input. We conducted the experiment using a random antibody sequence with a defined region of interest (ROI) of 20 amino acids. Single-site and multi-site mutations are performed within the ROI, followed by affinity computation (screening) of the resulting mutation space. Table 3 demonstrates the computational time cost on different sizes of virtual libraries using sequenced-based SentinusAI and structured-based approaches.

**Table 3.** Computational Cost of Antibody Maturation Space Search (sequence length = 20 amino acids)

| Methods | Input | Single-site Mutations | Double-site Mutations | Three-site Mutations | Four-site Mutations |
| --- | --- | --- | --- | --- | --- |
| | Mutation space | $10^2$ | $10^4$ | $10^6$ | $10^8$ |
| Schrodinger<sup>[28]</sup> | Structure | 4 days | ~100 days | Not applicable | Not applicable |
| Prodigy<sup>[29]</sup> | Structure | 336 seconds | 17.7 hours | 89 days | Not applicable |
| SentinusAI | Sequence | < 2min | 6 min | 40 minutes | 3 days |

\*Not applicable: computational time >100 days

**Table 4:** List of CPE Assay Results

| Supplier Id | Main screen, IC50 (ng/ml) | Main screen, Activity status | Main screen, Max% inhibition |
| --- | --- | --- | --- |
| AIN2-1_REGN10933 | 1.21 | Active | 159.93 |
| AIN1-31_AINNL0031 | 2.704 | Active | 116.48 |
| AIN1-48_AINNL0048 | 14.076 | Active | 109.96 |
| AIN1-42_AINNL0042 | 36.134 | Active | 131.78 |
| AIN1-45_AINNL0045 | 42.743 | Active | 102.12 |
| AIN1-49_AINNL0049 | 89.647 | Active | 169.39 |
| AIN1-50_AINNL0050 | 89.943 | Active | 108.57 |
| AIN1-43_AINNL0043 | 196.18 | Active | 123.23 |
| AIN1-38_AINNL0038 | 199.241 | Active | 160.54 |
| AIN1-40_AINNL0040 | 1142.851 | Active | 138.69 |
| AIN1-47_AINNL0047 | 1880.585 | Active | 117.5 |

### Broad coronavirus neutralization

#### ELISA assay results

We selected the top 50 AI-designed antibody sequences (AINL1-AINL50) predicted to bind best to the RBD for synthesis and functional assessments in the first round (pre-Omicron). Indeed, most antibodies synthesized could bind to the RBD of the SARS-CoV-2 spike protein, usually reaching an oversaturated state (OD value was greater than 2.0) at the highest concentration tested. Some antibodies also bound well to the SARS-CoV-2 RBD protein at lower concentrations (0.03 μg/ml) (Figure 4). Both the first batch of 50 antibodies (AINL1-AINL50) and the second batch of 20 antibodies (AINL51-AINL70) are well-expressed and had a high hit rate for binding to B.1, Delta, and Omicron. The two batches yielded 14% and 40% triple cross-binding hit rates, respectively.

**Figure 4:**
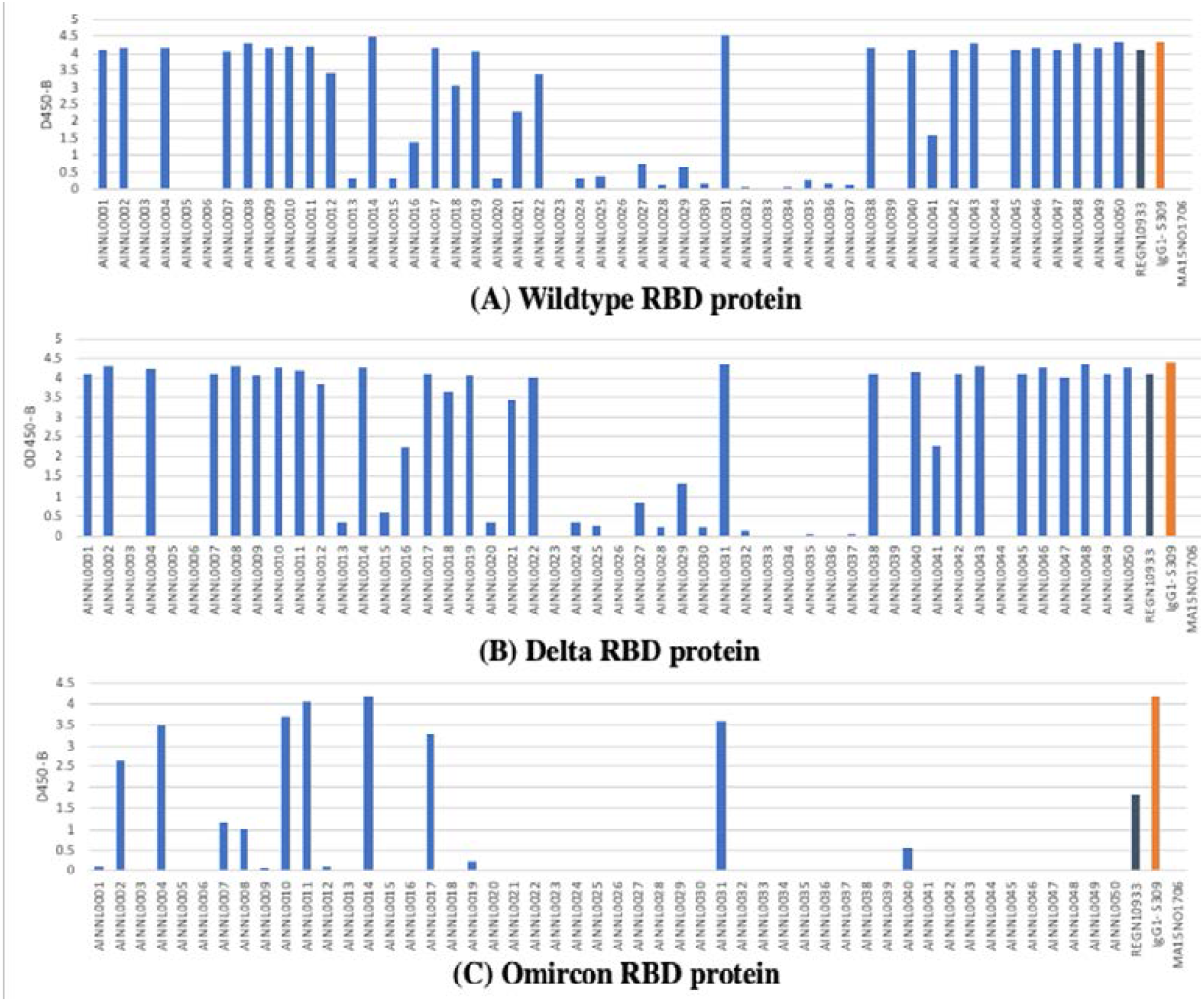
OD 450 absorbance values obtained by direct ELISA of 50 antibodies (1 μg/ml; blue) against wild type (A), delta (B) and Omicron (C). Plate coated with 1 ug/ml RBD protein. OD450 = optical density at 450 nm; Two therapeutic SARS-CoV-2 antibod-ies were tested as controls (black, orange) for variants.

We then measured the fold-change improvement in binding affinity of the designed antibody versus the template antibody as a measure of affinity maturation performance (Figure 5). Qualitative differences in antibodies are reflected by the shape of dose-response curves for antibody binding (Figure 6).

**Figure 5:**
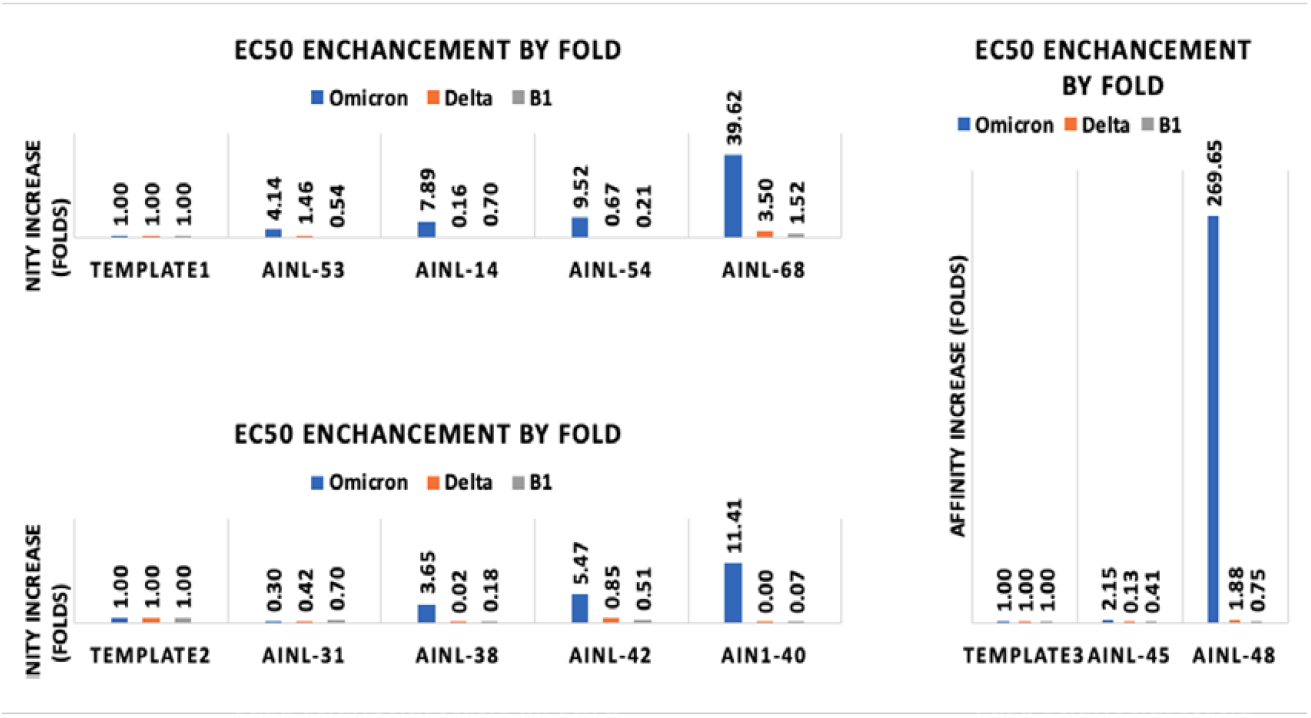
Affinity changes in designed antibodies compared to their template antibodies. (A) Template 1 (CR3022) and associated AI-designed antibodies, AINL53, AINL14, AINL54, AINL 68 (B) Template 2 (Regen 10933) and associated AI-designed antibodies, AINL31, AINL38, AINL42, AINL 40 (C) Template 3 (Regen 10987) and associated AI-designed antibodies, AINL45, AINL48.

**Figure 6:**
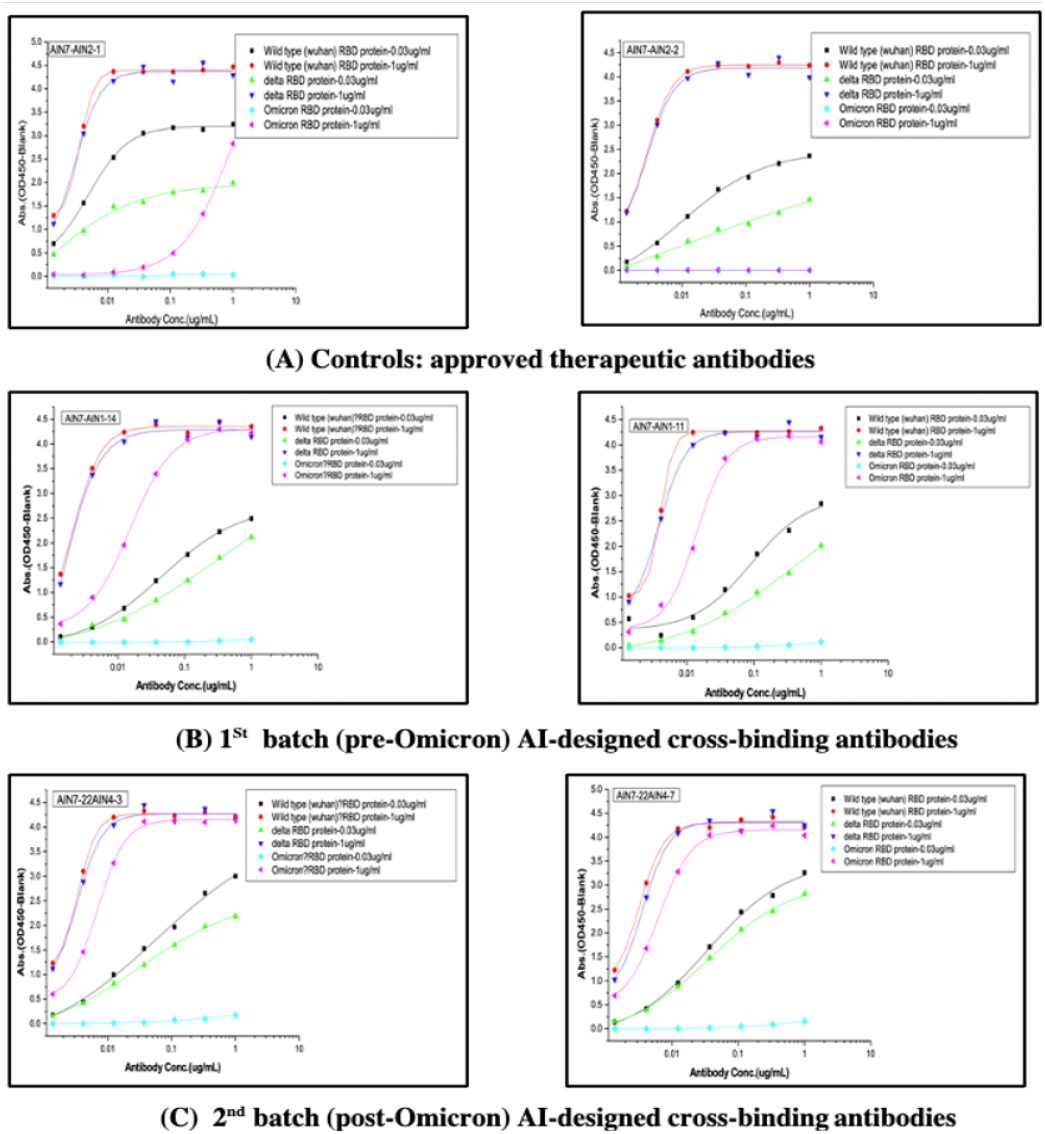
Selected ELISA curves for mAb binding to two different concentrations (0.03 µg/ml, 1 µg/ml) of RBD (WT, Delta, Omicron). (A) Controls of previously approved therapeutic antibodies (B) 1st batch AI-designed antibodies before Omicron prevalence (C) 2nd batch AI-designed antibodies after Omicron prevalence.

#### Pseudovirus-based neutralization assay

We next tested the ability of these antibodies to block pseudovirus entry into mammalian cells (Supplementary Data., Table S1), measured across different antibody concentrations and two strains (AINNL0001-AINNL0050 are neutralization results of the delta mutant pseudovirus, and AINNL0051-AINNL0070 are results from the omicron mutant pseudovirus). We conducted IC50 calculations only for antibodies showing inhibition rates of greater than 50%.

The AINNL0011 antibody inhibited pseudovirus entry by 91.98% at 20 μg/ml, with an IC50 of 8.03 μg/ml, indicating that the AINNL0011 antibody had a high neutralizing activity.

#### Coronavirus cytopathic assay

We next measured the ability of the designed antibodies to reduce the cytopathic effect (CPE) of the Delta and Omicron strains infecting Vero E6 host cells (Figure 7). Ten antibodies neutralized CPE of the Delta Strain (with IC50 <10 ug/ml), and one antibody blocked CPE for Omicron (Fig 6 and Table 5). For example, the IC50 of AINNL0031 was 2.704 ug/mL, indicating potent blockade of CPE. Overall, none of the antibodies showed significant direct toxicity (Supplementary Data. Table S2).

**Figure 7:**
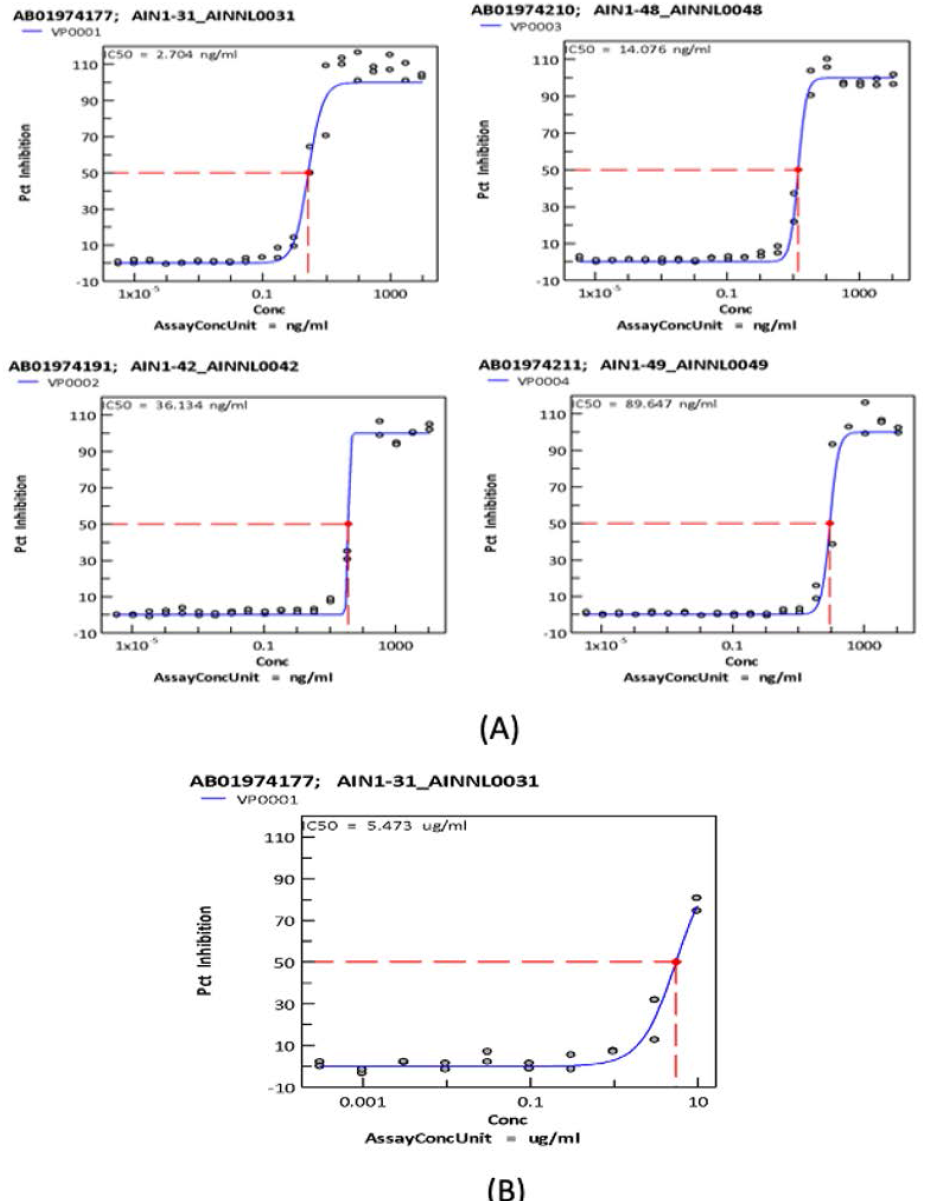
CPE inhibition curves of selected antibodies against Delta (A) and Omicron (B). Neutralization of the virus is done by mixing a fixed number of infectious virus particles with serial dilutions of the antibody and then adding this mixture to Vera E6 cells. Luminescence is read using a Perkin Elmer Envision or BMG CLARIOstar plate reader following incubation at room temperature for 10 minutes to measure cell viability.

## Discussion

Here, we tested the ability of an AI-based, structure-free approach to design antibodies with in vitro efficacy and at drastically reduced time and cost compared with traditional antibody engineering and affinity maturation strategies. This structure-free approach is critical, as current sequencing technologies have increased the number of protein sequences, but crystal structure generation is still a complex, risky, and time-consuming endeavor. Here we used an *in silico* antibody discovery approach based on affinity prediction models and deep learning techniques that incorporated only complex sequence information and no structural information. This modeling strategy captures functionally critical features encoded at the amino acid level that contribute to the interactions and the resulting binding affinities between antibody and antigen. This modeling at the amino acid level, rather than the atomic level, significantly reduces the time needed for both training and prediction and can also accommodate smaller annotated training datasets due to the model’s simplicity. This model showed improved accuracy compared to traditional methods as shown in Table 2 (A) and had comparable reliability shown in the same table for the benchmark study of traditional structure-based approaches on SKEMPI and AB-Bind datasets. The results from the SARS-COV2 RBD work (Figure 5) showed an affinity improvement of 11.41-, 39.62-, and 269.65-fold change with respect to each original antibody template, even though the antigens were not part of the training set, demonstrating the model’s utility even for novel antigens.

One interesting finding of our study was the discrepancy between the antibody binding data obtained through ELISA experiments and the observed neutralizing capacity of the antibody for blocking infection (either of the pseudovirus or in the CPE assays). This difference may arise from a number of factors, including differences in RBD structure when plate-bound in the ELISA assay versus its structure in the context of a live virus or the presence of different antibody binding sites on the RBD (e.g., antibodies have high-affinity interactions that may not physically block the RBD and receptor interaction). The ability to qualify antibody efficacy via functional assays remains an important validation step in therapeutic antibody development.

We also showed that our AI model could design cross-binding antibodies against many different antigen populations, including viral mutant strains. We hypothesized that the high dimensional features learned during the AI training process may represent components of the viral evolutionary process, and therefore could predict binding affinity in the virtual screening process even when it only contained known virus strains. This approach worked here for SARS-COV-2, where 14% of the 50 screened antibodies generated prior to Omicron still bound all the strains (omicron, delta and BA.1). Obtaining cross-reactivity for antibodies can indicate a lower specificity [31], but a broadly cross-reactive antibody (e.g., generated through engineering approaches [30]) may have better therapeutic potential in coping with viral evolution.

We also present a feasible computational workflow that allows for iterations with wet-lab results for better antibody design and cross-validation. This interaction is evident by the 40% cross-binding hit rate obtained in the second round of antibody synthesis (of 20 sequences), a significant performance leap compared to the 14% cross-binding hit rate in the first round.

This study demonstrated a highly cost efficient and effective approach towards the generation of therapeutic antibodies for single or multiple viral strains. However, an important consideration in the evaluation of this study’s effectiveness is that the neutralizing capability of the designed antibody lacks consistency with the binding affinity results. Multiple mechanisms and processes are involved in the neutralizing effects besides binding affinity. For example, the binding epitope on the spike protein and antibody conformation also impact the infection and translocation process during viral invasion. Therefore, epitope mapping and conformation dynamics studies are also required for a more precise design of neutralizing antibodies. In addition, in vivo efficacy studies were not performed yet, which limited the scope of this research.

Altogether, this study presents a novel method and model for both the initial design of therapeutic antibodies, as well as the ability to iteratively improve upon those initial designs to accommodate persistent mutations in the target protein of a rapidly evolving pathogenic virus. This combination of flexibility and throughput, while maintaining low computational burden, could be beneficial in other applications of the technology as well.

## Acknowledgements

This project is a collaborative effort of Ainnocence, Sino Biological, and Southern Research. We would like to give special thanks to the following people: We thank Cong Chen from Sino Biological for his help on ELISA and Pseudovirus-based neutralization assay. We thank Paige and Lynn Rasmussen from Southern Research for their help on CPE experiments.

We thank Junfeng Wu, Pinyu Xiao, and Vincent Brand for their engineering effort on the SentinusAI platform. We thank Sheng Ding, Iwan Alexander, Murat Tanik, and Mark Suto for their scientific advisory support on Ainnocence’s research works. We thank Raymond Price for his assistance with the manuscript.

The abovementioned 70 antibody sequences are filed in the application for Patent Cooperation Treaty (PCT) (Application id: PCT/CNT2022/094029)

