## Supplemental Files for "AI-Designed, Mutation-Resistant Broad Neutralizing Antibodies Against Multiple SARS-CoV-2 Strains"

### **Supplementary Data :**

Table S1: Results of Pseudovirus-based Neutralization Assay

| **Project ID** | **Sequence name** | **Conc.**（**μg/ml)-Inhibition rate**（**%**） | | | | | | **IC50** |
| --- | --- | --- | --- | --- | --- | --- | --- | --- |
|  |  | **100** | **20** | **4** | **0.8** | **0.16** | **0.032** |  |
| AIN1-11 | AINNL0011 | 75.60% | 91.98% | 17.96% | 30.31% | 14.00% | -14.42% | 8.03 |
| AIN2-1 | REGN10933 antibody, Human IgG1 mAb | 69.88% | 68.01% | 38.87% | 26.42% | 12.95% | 38.71% | 7.4 |
| TJ41A-2 | IgG1- S309 | 24.19% | 15.75% | 42.78% | 57.57% | 49.51% |  |  |
| Positive control | #N/A | 99.98% | 98.63% | 78.92% | 49.78% | 26.49% | 7.76% | 0.81 |

Table S2: List of Toxicity Assay Results

| **Supplier Id** | **TOX,**  **CC50**  **(ng/mL)** | **TOX,**  **Activity**  **Status** | **TOX,**  **Min %**  **Viability** | **TOX,**  **Max %**  **Viability** |
| --- | --- | --- | --- | --- |
| AIN2-1_REGN10933 | >10000 | Inactive | 90.13 | 98.84 |
| AIN1-31_AINNL0031 | >9780 | Inactive | 91.49 | 104.54 |
| AIN1-48_AINNL0048 | >11000 | Inactive | 91.20 | 98.83 |
| AIN1-42_AINNL0042 | >10890 | Inactive | 91.09 | 108.57 |
| AIN1-45_AINNL0045 | >11440 | Inactive | 89.69 | 102.73 |
| AIN1-49_AINNL0049 | >11220 | Inactive | 93.35 | 103.61 |
| AIN1-50_AINNL0050 | >11220 | Inactive | 94.01 | 103.53 |
| AIN1-43_AINNL0043 | >12330 | Inactive | 91.59 | 101.78 |
| AIN1-38_AINNL0038 | >10890 | Inactive | 95.64 | 101.41 |
| AIN1-40_AINNL0040 | >10780 | Inactive | 88.33 | 102.83 |
| AIN1-47_AINNL0047 | >24780 | Inactive | 90.25 | 104.14 |
